# Increasing phagocytosis of microglia through targeting CD33 with liposomes displaying glycan ligands

**DOI:** 10.1101/2021.05.08.443135

**Authors:** Abhishek Bhattacherjee, Gour Chand Daskhan, Arjun Bains, Adrianne E. S. Watson, Ghazaleh Eskandari-Sedighi, Chris D. St. Laurent, Anastassia Voronova, Matthew S. Macauley

## Abstract

CD33 is an immunomodulatory receptor expressed on microglia and genetically linked to Alzheimer’s disease (AD) susceptibility. While antibodies targeting CD33 have entered clinical trials to treat neurodegeneration, it is unknown whether the glycan-binding properties of CD33 can be exploited to modulate microglia. Here, we use liposomes that multivalently display glycan ligands of CD33 (CD33L liposomes) to engage CD33. We find that CD33L liposomes increase phagocytosis of cultured monocytic cells and microglia in a CD33-dependent manner. Enhanced phagocytosis strongly correlates with loss of CD33 from the cell surface and internalization of liposomes. Increased phagocytosis by treatment with CD33L liposomes is dependent on a key intracellular signaling motif on CD33 as well as the glycan-binding ability of CD33. These effects are specific to *trans* engagement of CD33 by CD33L liposomes, as *cis* engagement through insertion of lipid-linked CD33L into cells produces the opposite effect on phagocytosis. Moreover, intracerebroventricular injection of CD33L liposomes in mice enhances phagocytosis of microglia in a CD33-dependent manner. These results demonstrate that multivalent engagement of CD33 with glycan ligands can modulate microglial cell function.

## Introduction

Genes correlating with Alzheimer’s disease (AD) susceptibility converge on immunoregulation, particularly those expressed in microglia.^1^ As the major immune cell in the brain, microglia play a myriad of roles and are connected with AD progression through their ability to phagocytose amyloid-β.^2^ One of the key genes linked to AD susceptibility is the glycan-binding protein, CD33.^3^ The link between CD33 and AD susceptibility was identified through genomewide association studies, where a rare allele of CD33 is AD protective.^4,5^ This rare allele undergoes alternative mRNA splicing,^6^ leading to reduced expression of a long isoform of CD33 (CD33M; M=Major isoform), and increased expression of a short isoform (CD33m; m=minor isoform). Understanding the individual roles of these two CD33 isoforms has been challenging since human cells express a mixture of these two CD33 isoforms. Moreover, murine CD33 has several highly diverget features and does not undergo alternative mRNA splicing.^7^ Nevertheless, significant progress has recently been made and supports a role for CD33M in repressing phagocytosis, while distinct roles for CD33m have been also identified recently.^7–10^

Therapeutic targeting of proteins implicated in AD susceptibility has become an attractive approach. For example, antibodies targeting TREM2 have undergone rigorous evaluation in mouse models,^11–13^ and are now under clinical evaluation (NCT03635047). These antagonistic anti-TREM2 antibodies stimulate microglial cell proliferation and phagocytosis as a key means of skewing microglia into a disease protective phenotype.^13^ TREM2 is proposed to be negatively regulated by CD33,^14,15^ and antibodies targeting CD33 – which have a long history in the clinic for the treatment of leukemia^16^ – are being assessed as means of modulating microglia in a clinical trial (NCT03822208). Anti-CD33 antibodies deplete CD33M from the surface,^17–20^ which is where it needs to be in order to repress phagocytosis.^8^ However, as murine CD33 does not serve as a good model for human CD33,^7^ *in vivo* pre-clinical testing of anti-CD33 antibodies have not been carried out.

CD33 is a member of sialic acid-binding immunoglobulin-type lectin (Siglec) family of immunomodulatory receptors.^21^ CD33 binds to both α2-3 and α2-6 linked sialosides, with emerging evidence that binding may be enhanced by underlying sulfation.^22,23^ Engaging Siglecs with high-affinity and specific glycan ligands is an alternative approach compared to antibodies.^24–28^ A high-affinity selective glycan ligand for targeting human CD33 was previously developed through dual modification of the 5- and 9-position of sialic acid with hydrophobic substituents.^29^ Recently, a high-resolution crystal structure of CD33 and this high-affinity CD33 ligand was determined, providing a basis for how the modifications exploit surrounding hydrophobic pockets.^30^ Moreover, this same study found that microparticles displaying this CD33 ligand increased the uptake of Aβ peptide in human monocytic cells (THP1), although the mechanism by which this occurred was not elucidated. Regardless, these results suggest that targeting CD33 through its glycan-binding domain has the potential to modulate microglial phagocytosis.

Here, we developed and implemented liposomes displaying the high-affinity CD33 ligand (CD33L) for engaging CD33 and show that CD33L liposomes enhance phagocytosis in a CD33-dependent manner. CD33L liposomes were formulated with two different fluorophores – one for monitoring CD33-engagement and a pH-sensitive fluorophore for monitoring cellular internalization – that led to the discovery that the ability of CD33L liposomes to enhance phagocytosis strongly correlates with CD33 and liposome internalization within the cells. Moreover, CD33L liposomes enhance phagocytosis of primary mouse microglia expressing human CD33 following an intracerebroventricular injection. These results demonstrate the pharmacological potential of modulating immune cell function in the brain through targeting an immunomodulatory glycan-binding protein.

## Results and Discussion

### Targeting CD33 with CD33L liposomes

A bifunctionally-modified version of Neu5Acα2-6Galβ1-4Glc (α2-3 sialyl-lactose) **1** that has both selectivity^29^ and increased affinity (*K*_d_ = 87 μM)^22^ for CD33 was used in these studies. For incorporation into liposomal nanoparticles, this synthetic CD33 ligand (CD33L) was conjugated to peggylated distearoylphosphatidylethanolamine (PEG-DSPE) to form CD33L-PEG-DSPE **3** (**Fig. 1a**). For monitoring binding and internalization of liposomes to CD33-expressing cells, two fluorophores - AlexaFluor647 (AF647) and pH-sensitive rhodamine (pHrodo), respectively – were also conjugated to lipid to form AF647-PEG-DSPE **7** (**Fig. 1b**) and pHrodo-PEG-DSPE **6** (**Fig. 1c**), respectively. With these lipid conjugates in hand, liposomal nanoparticles were formulated to contain 0.1 mol % AF647-PEG-DSPE **7**, 0.1 mol % pHrodo-PEG-DSPE **6**, and with or without 3 mol% CD33L-PEG-DSPE **3** to make naked liposomes (**Fig. 1d**) and CD33L liposomes (**Fig. 1e**), respectively. The liposome formulation was modeled on an FDA-approved formulation that is well tolerated *in vivo*.^31–33^ Specifically, the base formulation of these liposomes contained a total of 5 mol % PEG-DSPE, along with 57% distearoylphosphatidylcholine, 38% cholesterol, and were extruded to 120 +/-20 nm in size as determined by dynamic light scattering.

**Fig. 1.**
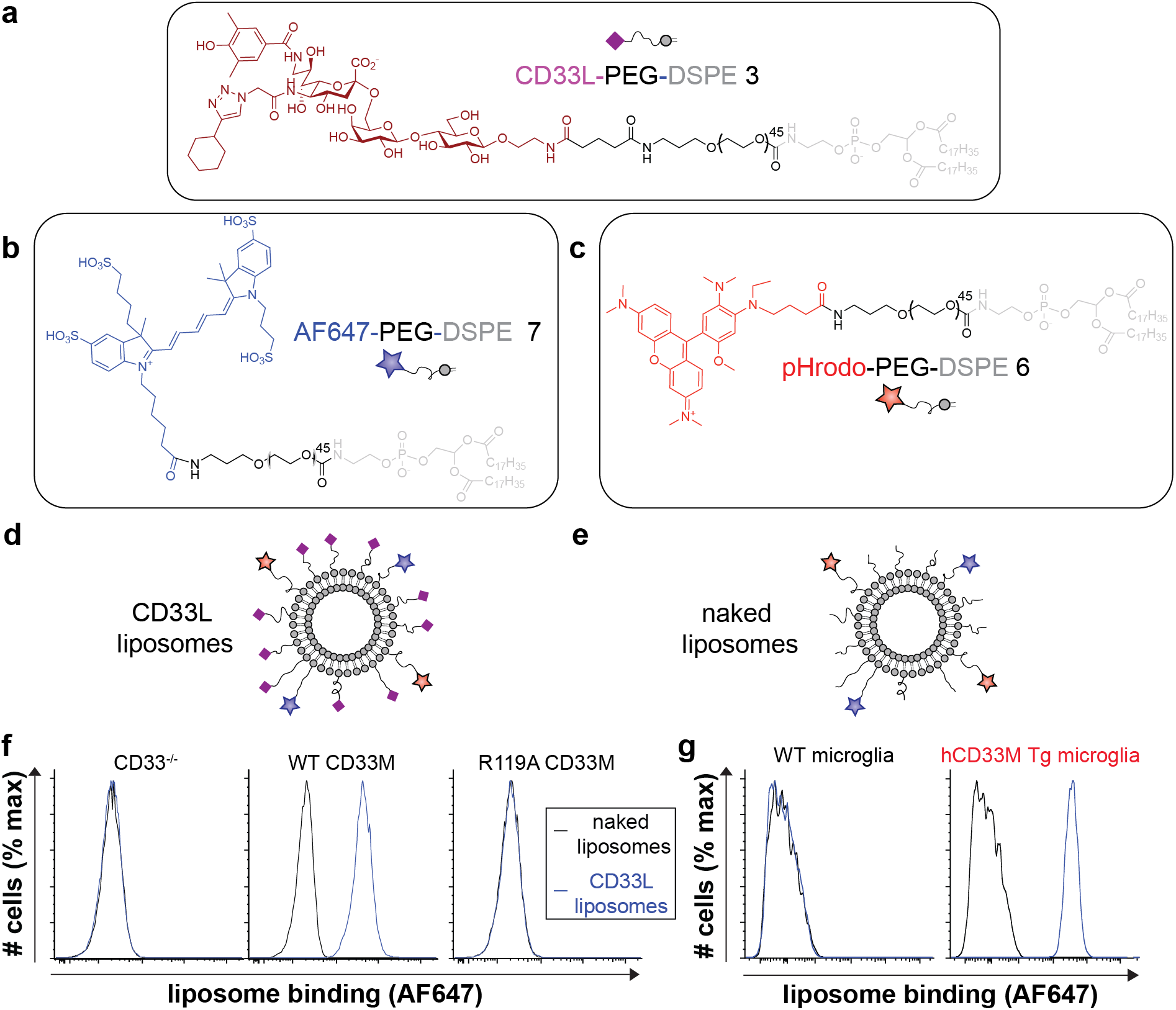
Fluorophore conjugated CD33L liposome formulation for engaging CD33 on cells. (**a-c**) Chemical structures of CD33L-PEG-DSPE **3** (**a**), AF647-PEG-DSPE **7** (**b**), and pHrodo-PEG-DSPE **6** (**c**). (**d,e**) Cartoon diagrams of CD33L (**d**) and naked (**e**) liposomes. (**f**) Binding of naked (black) and CD33L liposomes (blue) to CD33^-/-^ (*middle panel*), and CD33^-/-^ U937 cells transduced with WT CD33M (*middle panel*) or R119A CD33M (*right panel*). (**g**) Binding of naked (black) and CD33L liposomes (blue) to primary microglia isolated from WT (*Cx_3_cre^Cre-/+^hCD33M^-/-^*) or hCD33M transgenic (Tg; *Cx_3_cre^Cre-/-^hCD33M^-/-^*) mice. Binding of liposomes to cells was measured by total AF647 signal by flow cytometry.

Liposomes were initially tested for binding to U937 cells by flow cytometry using the AF647 signal as a measure of total binding. CD33L liposomes showed robust binding to WT U937 cells, with no binding to CD33^-/-^ U937 cells or CD33^-/-^ U937 cells transduced with an R119A mutant of CD33 (**Fig. 1f**). Likewise, CD33L liposomes bound strongly to primary mouse microglia from human CD33M transgenic mice but not WT mouse microglia (**Fig. 1g**). These results demonstrate that CD33L liposomes specifically engage hCD33M on the cell surface of immune cells, which is in line with previous observations using a multivalent display of this ligand.^29,32^

### CD33L liposomes increase phagocytosis in a CD33-dependent manner

We examined the effect of engaging CD33 with CD33L liposomes on phagocytosis by incubating cells with liposomes, washing the cells, and monitoring phagocytosis using fluorescent carboxylate-modified polystyrene beads in conjunction with flow cytometry. Initial experiments with the standard formulation (3.3 mol % CD33L liposomes) significantly enhanced phagocytosis in WT but not CD33^-/-^ U937 cells, therefore, a series of CD33L liposomes were prepared with decreasing amounts of CD33L (**Fig. 2a,b**). Effects on phagocytosis were lost at CD33L densities lower than 3.3 mol %. Therefore, we moved forward with 3.3 mol % CD33L density for further optimizations. Varying the concentration of CD33L liposomes, we find that enhanced phagocytosis was strongly correlated with the concentration of CD33L liposomes used and, notably, the highest concentration increased phagocytosis of WT U937 cells to nearly the level of CD33^-/-^ cells (**Fig. 2c,d**). As a final optimization, 3.3 mol % CD33L liposomes at a concentration of 100 μM were incubated for different amounts of time, with results indicating that phagocytosis increased for up to 2 hr but no further increase was observed beyond 2 hr (**Fig. 2e,f**). This data indicates that engagement of CD33 by CD33L liposomes increases phagocytosis in U937 cells in a manner that is dependent on CD33L density, liposome concentration, and time with which the liposomes are incubated with cells.

**Fig. 2.**
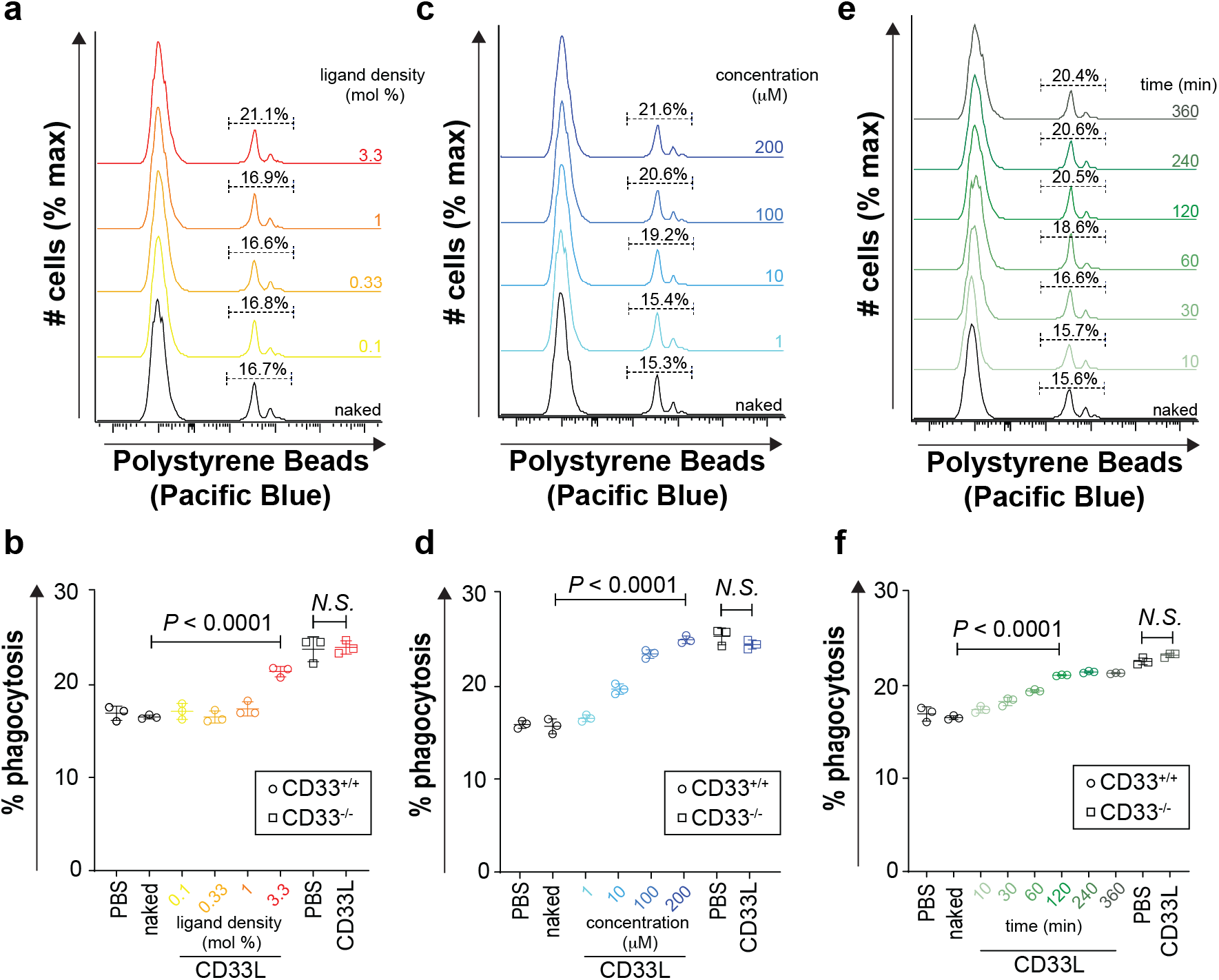
CD33L liposomes increase phagocytosis of polystyrene beads in U937 cells. (**a,b**) Liposomal nanoparticles were formulated at different CD33L densities and tested for their ability to modulate phagocytosis. (**c,d**) The most efficient formulation (3.33 mol % CD33L) was selected and tested at different liposomal concentrations. (**e,f**) Liposomal concertation of 100 μM was selected and incubated with cells for various times before examining phagocytosis. Phagocytosis was measured by flow cytometry (**a,c,e**) and the percentage of phagocytic cells was quantified (**b,d,f**). Statistical significance was calculated using a one-way ANOVA with Dunnett’s test.

Recently, we and others have demonstrated that the CD33 long isoform (CD33M) represses microglial phagocytosis, which is consistent with CD33M acting as an immunoinhibitory receptor.^7–10^ These results suggest that engaging CD33 with CD33L liposomes reverses this inhibitory effect. This is conceptually similar to results observed for another member of the Siglec family, CD22, where anti-CD22 antibodies enhance microglial phagocytosis.^34^

### CD33L liposomes deplete cell surface CD33

As Siglecs are endocytic receptors,^35^ we investigated whether CD33L liposomes induce internalization of CD33. Internalization was assessed by depletion of CD33 from the cell surface using cell surface staining with an anti-CD33 antibody (clone HIM3-4)^36^ that does not bind the glycan-binding domain of CD33. In parallel, internalization of the liposomes was assessed using the fluorescent signal from pHrodo incorporated into the liposomes and comparing it to the fluorescent signal from AF647 that represents total liposome binding. For liposomes where the CD33L density was varied, 3.3 mol % CD33L induced the most significant depletion of CD33 from the cell surface (50% depletion), liposome internalization, and liposome binding (**Fig. 3a-f**). Liposomes bearing 1 mol % CD33L produced a modest depletion of cell surface CD33 levels as well as CD33 internalization, yet did not increase phagocytosis (**Fig. 2a,b**). Examining the concentration-(**Fig. 3g-l**) and time-dependent (**Fig. 3m-r**) effects of CD33L liposomes on internalization revealed that internalization of both CD33 and the CD33L liposomes strongly correlate with enhanced phagocytosis. Specifically, 200 μM liposomes led to the greatest internalization of CD33 and liposomes, while internalization plateaued at 2 hr.

**Fig. 3.**
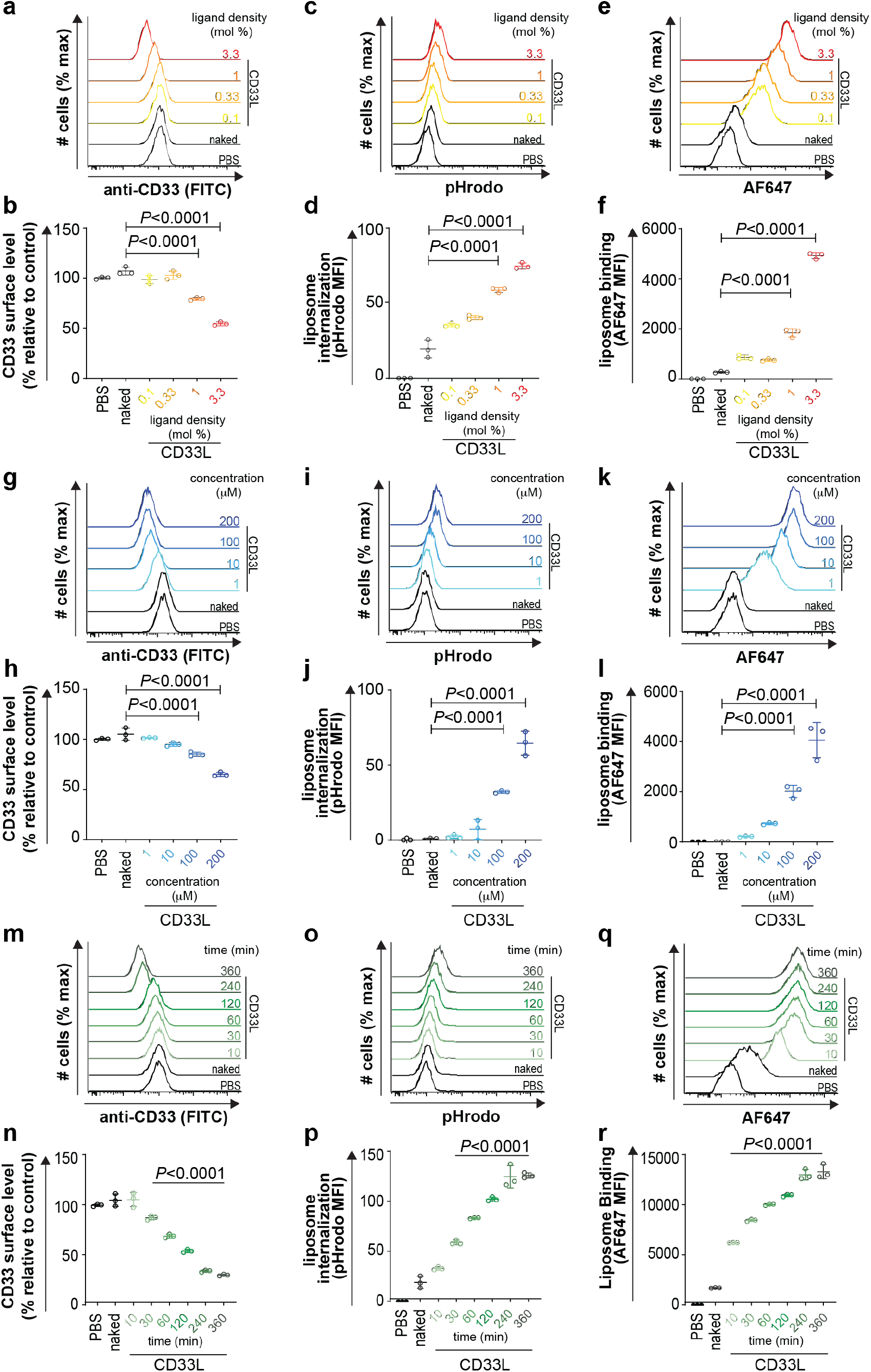
CD33L liposomes induce internalization of CD33. (**a-f**) CD33L liposomes formulated with different CD33L densities, containing pHrodo and Alexa 647, were incubated with WT U937 cells. (**g-l**) CD33L liposomes formulated with 3.3% CD33L were incubated with WT U937 cells at different concentrations. (**m-r**) CD33L liposomes were formulated with 3.3% CD33L and incubated with U937 cells at 100 μM for different amounts of time. Cell surface CD33M cell surface staining (**a,b,g,h,m,n**), internalized liposomes (**c,d,i,j,o,p**), and total liposome binding (**e,f,k,l,q,r**) measured by flow cytometry. Raw flow cytometry plots (**a,c,e,g,i,k,m,o,q**) and summary plots of the MFI values (**b,d,f,h,j,l,n,p,r**) are colour coded for the conditions used. Assays were replicated three times and statistical significance was calculated based on a one-way ANOVA with Dunnett’s test.

We also examined CD33 and liposome internalization by confocal microscopy. Following incubation of U937 cells with CD33L liposomes, a 46% decrease in cell surface CD33 was observed (**Supplementary Figure S1a,b**), which are highly consistent with the flow cytometry results. In these images, pHrodo and AF647 signal from the liposomes were also readily apparent in most CD33-expressing cells, where this signal was largely absent in cells incubated with naked liposomes. Using antibody HIM3-4 to track the CD33 inside the cell was not successful, therefore, a C-terminal 3xFLAG tagged version of CD33 was expressed in CD33^-/-^ U937 cells. This approach enabled fluorescence tracking of CD33, using an anti-FLAG antibody, where a strong signal was observed around the pHrodo and AF647 signal (**Supplementary Figure S1c**). These results strongly suggest that CD33 is internalized along with the CD33L liposomes.

Previous work using 5 μM beads displaying this same ligand also induced an increase in phagocytosis, but due to the very large size of these beads it is unknown if they can be internalized and whether depletion of cell surface CD33 was at play.^30^ Our results demonstrate that internalization of CD33 closely correlates with increases in phagocytosis induced by CD33L liposomes. Interestingly, total CD33 engagement did not necessarily correlate with internalization and, consequently, increased phagocytosis. For example. CD33L liposomes bearing less than 3.3 mol % CD33L still bound to CD33-expressing cells as evidenced by the AF647 signal (**Fig. 3e**), but did not deplete CD33 from the cell surface (**Fig. 3a**) or increase phagocytosis (**Fig. 2a**).

Contrasting with this result,10 μM of liposomes (with the optimal 3.3 mol % CD33L), had a similar overall ability to bind to cells (**Fig. 3k**) but did deplete cell surface CD33 levels (**Fig. 3g**) and increase phagocytosis (**Fig. 2c**). Therefore, the degree of crosslinking of CD33 on the cell surface, which is dictated by both the density and affinity of the CD33L, is the most critical element dictating internalization of CD33.

### Role of glycan-binding and signaling motifs on phagocytosis

Binding of CD33 to its glycan ligands and control of immune cell signaling is dependent on a key salt bridge formed between Arg119 within its N-terminal V-set domain^30^ and signaling motifs within its cytoplasmic tail at Tyr340 (ITIM) and Tyr358 (ITIM-like)^37^, respectively. CD33^-/-^ U937 cells were previously established with WT CD33, R119A CD33, Y340A CD33, and Y358A CD33 re-introduced into these cells through lentiviral transduction.^8^ Here, these mutants were used to interrogate the roles of these key residues in the enhancement of phagocytosis by CD33L liposomes. Cell lines were treated with 100 μM CD33L liposomes (3.3 mol% CD33L) for 60 min before carrying out a phagocytosis assay with polystyrene beads. Compared to CD33^-/-^ cells (**Fig 4a,b**), cells reconstituted with WT CD33 showed enhanced phagocytosis by the treatment of CD33L liposomes (**Fig. 4c,d**). In both the R119A (**Fig. 4e,f**) and Y340A (**Fig. 4g,h**) mutant, the phagocytosis-enhancing effect of CD33L liposomes was lost. CD33L liposomes were still capable of enhanced phagocytosis in cells expressing the Y358A mutant (**Fig. 4i,j**). Previously, we showed that Y340 was critical for the ability of CD33 to repress phagocytosis, and others have also shown Y340 is required for antibody-mediated endocytosis of CD33.^19,20^ Consistent with these previous results, we find that the Y340A CD33 mutant internalizes significantly less as compared to WT CD33 in response to CD33L liposomes, whereas the Y358A CD33 mutant had no defect in internalization (**Fig. 4K**).

**Fig. 4.**
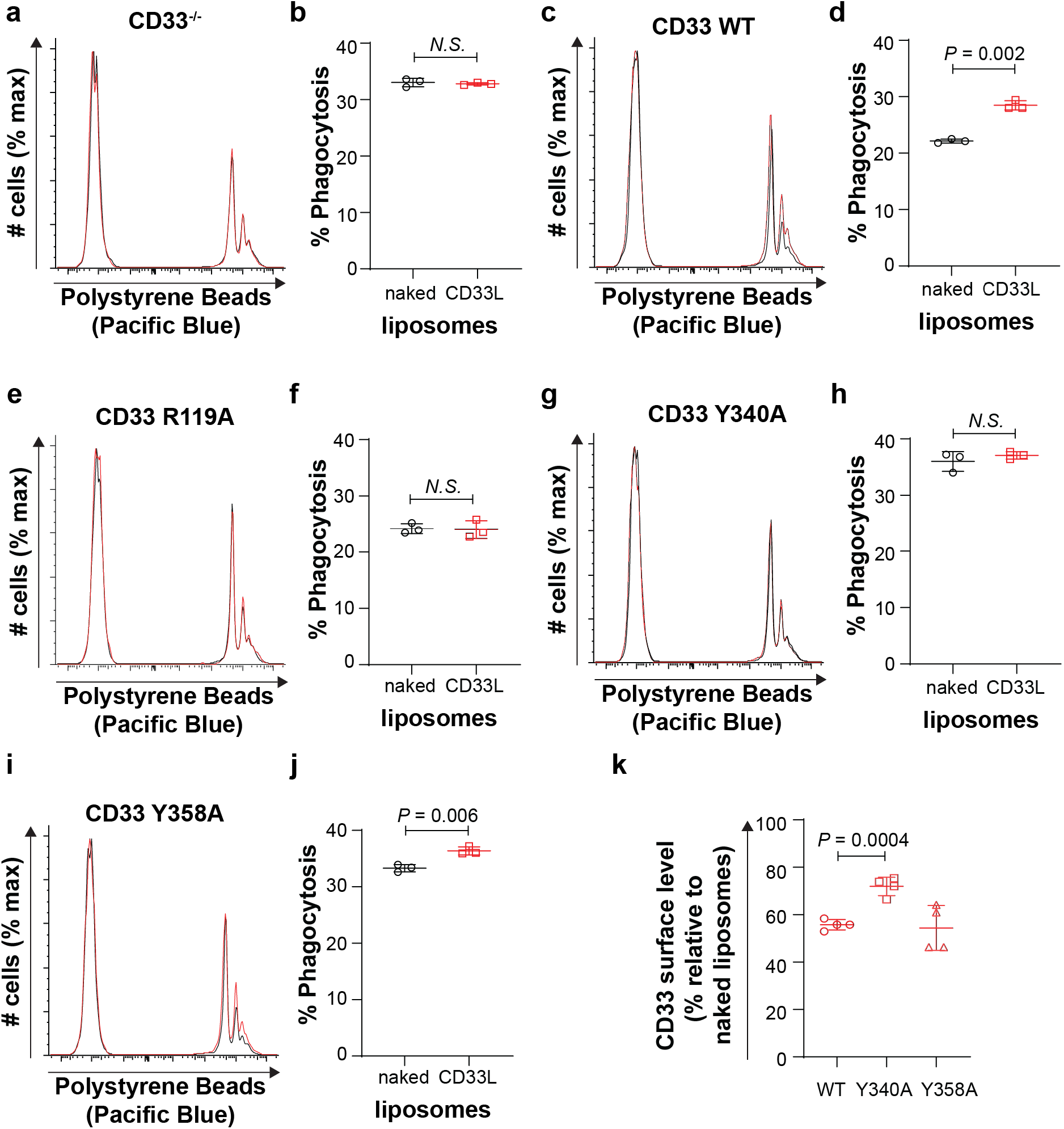
Increased phagocytosis by CD33L liposomes is dependent on the glycan-binding and signaling abilities of CD33. Naked (black) or CD33L (red) liposomes were incubated with CD33^-/-^ U937 cells transduced with (**a,b**) empty vector, (**c,d**) WT CD33M, (**e,f**) R119A CD33M, (**g,h**) Y340A CD33M, (**i,j**) and Y358A CD33M prior to performing phagocytosis assay. Raw flow cytometry plots (**a,c,e,g,i**) and summary plots of the MFI values (**b,d,f,h,j**) are shown for each cell type. (**k**) CD33M internalization following incubation with naked liposomes or CD33L liposomes was examined on WT and CD33^-/-^ U937 cells transduced with Y340A or Y358A CD33M mutant. Assays were replicated three times and statistical significance was calculated based on a oneway ANOVA with Dunnett’s test.

These results establish that the ability of CD33 to bind its glycan ligands, repress signaling, and internalize in response to CD33L liposomes are essential for the ability of CD33L liposomes to enhance phagocytosis. Recently, we showed that the ITIM of CD33 is essential for its ability to repress phagocytosis, whereas the ITIM-like residue is dispensable.^8^ Our findings here demonstrate that crosslinking alone of CD33, without its ability to modulate immune cell signaling through the ITIM, is insufficient for CD33L liposomes to impact phagocytosis. Previous results demonstrating a requirement for the ITIM of CD33 in antibody-mediated internalization^20^ are in agreement with our data. The optimized formulation of CD33L liposomes bares somewhere on the order of 1000-2000 molecules of CD33L based on a 100 nm liposomes containing 84,000 lipids. This high multivalency along with our 120 nm liposomes falling within the range of efficient clathrin-mediated endocytosis gives these liposomes a strong ability to internalize surface receptors.^38–41^

### Engaging CD33 in cis has the opposite impact on phagocytosis

Siglecs can be engaged by ligands on other cells (*trans*) or the same cell that the Siglec is expressed (*cis*)^21^. Results above with liposomes represent engagement with *trans* ligands. To test the impact of engagement with *cis* ligands, CD33L-PEG-DSPE **3** was inserted into cells (**Supplementary Figure S2a**). Successful insertion of CD33L into U937 cells was demonstrated by enhanced staining with CD33-Fc (**Supplementary Figure S2b**). Unlike *trans* engagement, *cis* engagement of CD33 with inserted CD33L led to a significant decrease in phagocytosis (**Supplementary Figure S2c,d**) in U937 cells expressing WT CD33. Insertion of the CD33L-PEG-DSPE **3** had no impact on phagocytosis within CD33^-/-^ U937 cells (**Supplementary Figure S2e,f**), nor did it impact phagocytosis in CD33^-/-^ U937 cells expressing R119A CD33 (**Supplementary Figure S2g,h**) or Y340A CD33 (**Supplementary Figure S2i,j**). These results are similar to recent work in which high-affinity glycan ligands of Siglec-9 were inserted into cells, which induced a suppressive signal.^42^ The basis for how the engagement of CD33 with *cis* and *trans* ligands produce the opposite impact on phagocytosis is not completely understood, but results clearly indicate that the ITIM of CD33 is essential for both processes, which offers strong evidence for an immunomodulatory function of CD33.

### CD33L liposomes increase phagocytosis in primary microglia ex vivo

Results demonstrating that CD33L liposomes enhance phagocytosis in cultured U937 cells motivated us to test if they also have a similar effect in microglia. As mouse CD33 is not functionally conserved with human CD33 (hCD33), we previously developed a transgenic mouse model in which hCD33M is expressed in the microglial cell lineage and represses phagocytosis.^7^ We isolated microglia from WT and hCD33M transgenic mice and performed a competitive phagocytosis assay following incubation of cells with liposomes (**Fig. 5a**). Phagocytosis within the hCD33M^+^ and hCD33M^-^ populations were deconvoluted using the gating scheme shown in **Figure 5b**. The mixture of cells was incubated with either PBS (**Fig. 5c,d**), naked liposomes (**Fig. 5,e,f**), or CD33L liposomes (**Fig. 5g,h**). hCD33M transgenic microglia showed a decrease in phagocytosis compared to WT microglia in conditions where cells were treated with PBS and naked liposomes, which is anticipated based on the ability of hCD33M to repress phagocytosis in these transgenic mice^7,8^. Pre-treatment with CD33L liposome increased phagocytosis in the CD33M microglia (**Fig. 5i**), such that no statistically significant difference was observed in phagocytosis between the WT and hCD33M transgenic microglia. These results demonstrate that CD33L liposomes can enhance microglial cell phagocytosis in a CD33-dependent manner.

**Fig. 5.**
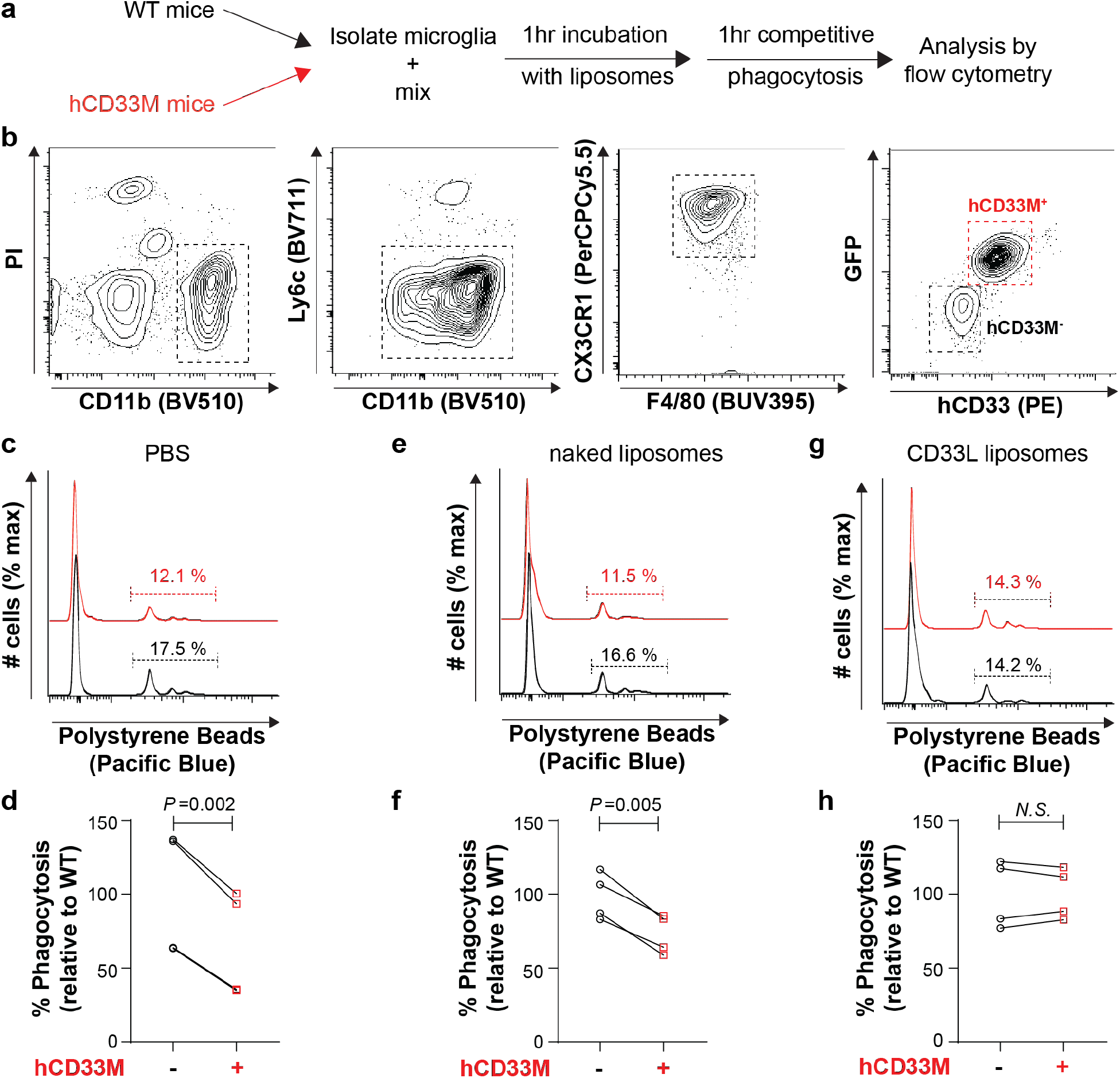
CD33L liposomes enhance phagocytosis in hCD33M transgenic mouse microglia. (**a**) Schematic for a competitive phagocytosis assay with primary microglia taken from WT (black) and hCD33M transgenic (red) mice. (**b**) Gating strategy for the competitive phagocytosis assay. (**c-h**) Results of the competitive flow cytometry-based uptake of polystyrene microbeads. Isolated microglia were pretreated with PBS (**c,d**), naked liposome (**e,f**), or CD33L liposomes (**g,h**). Raw flow cytometric data (**c,e,g**) and the percentage of phagocytosis was quantified in each condition (**d,f,h**). (**i**) Absolute phagocytosis values in the hCD33M Tg microglia in the three conditions. Each point represents an independent experiment from different mice. *N.S*. represents no statistical significance based on a paired Student’s t-test.

### In vivo administration of CD33L liposomes enhanced phagocytosis in microglia

An advantage of our liposome formulation is that it is based on an FDA-approved formulation and is well-tolerated *in vivo*.^32^ However, this formulation does not penetrate the blood-brain barrier.^43^ Therefore, to test whether CD33L liposomes can successfully stimulate microglial phagocytosis *in vivo*, we administered liposomes via ICV injection. Briefly, WT and hCD33M transgenic mice were ICV injected with PBS, naked liposomes, or CD33L liposomes into the right ventricle, followed by a 4 hr recovery prior to euthanization of mice. After euthanization brain samples were collected and microglia were isolated from each group (**Fig. 6a**). Initially, we performed a test with AF647 labeled naked and CD33L liposomes and examined AF647 fluorescence in isolated microglia, which demonstrated successful targeting of hCD33M-expressing cells selectively with CD33L liposomes (**Fig. 6b,c**). For assessing the impact on phagocytosis, WT or hCD33M Tg mice were carried out in parallel, and a competitive phagocytosis assay was used. In these studies, we assessed phagocytosis using the more physiologically-relevant aggregated fluorescent Aβ_1-42_ and used conditions with and without Cytochalasin-D to control for non-specific sticking or uptake. Consistent with our results described above, hCD33M microglia mice injected with PBS (**Fig. 6d,e**) or naked liposome (**Fig. 6f,g**) showed repressed phagocytosis compared to their WT counterparts. On the other hand, CD33L liposome injected hCD33M mice (**Fig. 6h,i**) showed a similar level of phagocytosis as compared to WT mice under identical conditions. This data demonstrates that *in vivo* administration of CD33L liposome can modulate phagocytosis in microglia in a CD33-dependent manner.

**Fig. 6.**
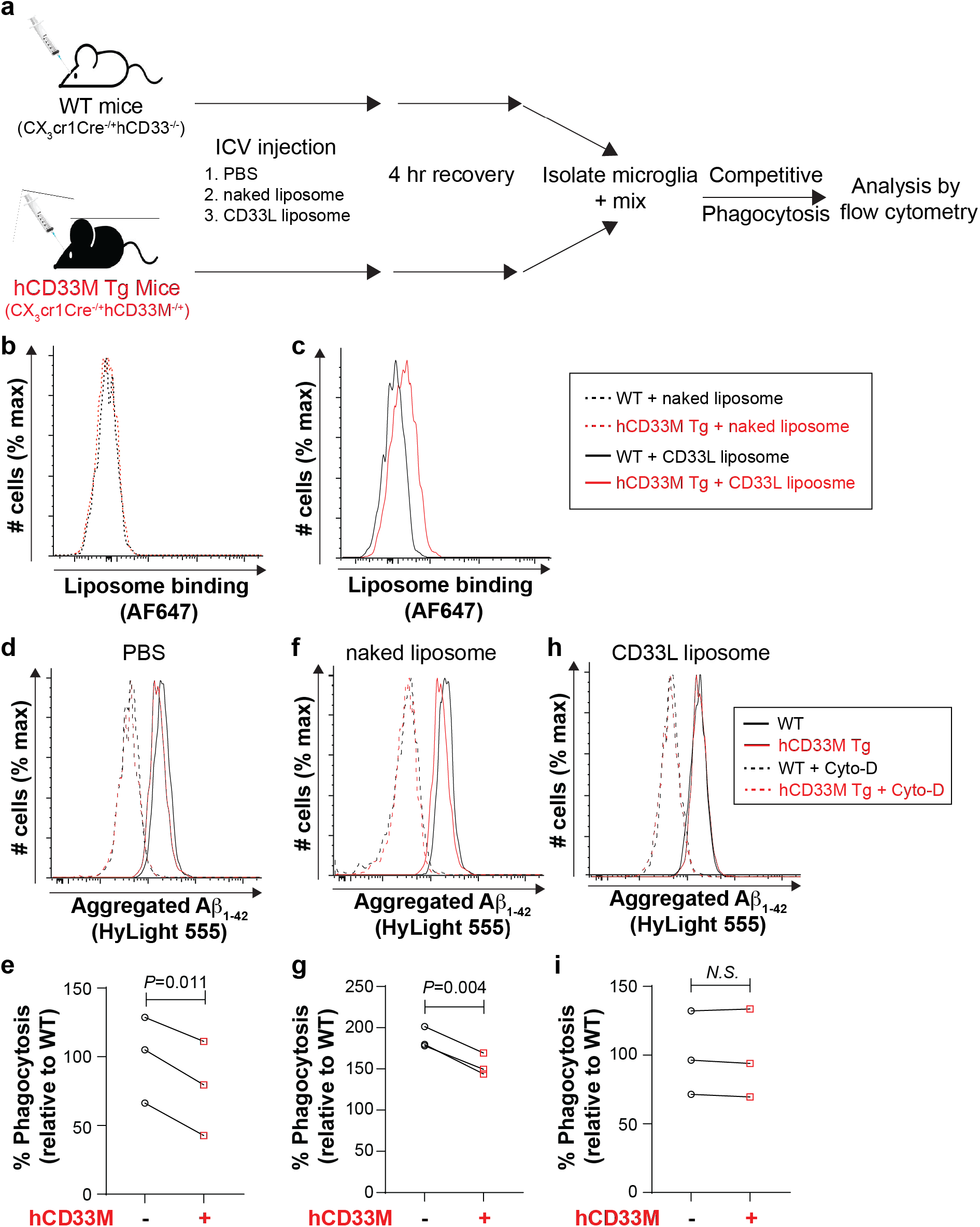
Intracerebroventricular injection (ICV) of CD33L liposomes modulate phagocytosis of hCD33M microglia. (**a**) Schematic representation of the experimental plan, in which WT (black) or hCD33M transgenic mice (red) were administered PBS, naked liposomes, or CD33L liposomes, followed by isolation and mixing of microglia, and a competitive phagocytosis assay. (**b,c**) AF647 signal in microglia after injection of naked or CD33L liposomes. (**d-i**) Results of the competitive flow cytometry-based uptake of aggregated Aβ_1-42_. WT (black) and hCD33M transgenic mice (red) in mice injected with PBS (**d,e**), naked liposome (**f,g**), or CD33L liposomes (**h,i**). Raw flow cytometric data (**d,f,h**) and the percentage of phagocytosis was quantified in each condition (**e,g,i**). Each point represents an independent experiment from different mice. *N.S*. represents no statistical significance based on a paired Student’s t-test.

In summary, our results demonstrate that CD33L liposomes can increase microglial phagocytosis both *in vitro* and *in vivo*. CD33L conjugated to microparticles produced a similar impact on phagocytosis *in vitro*, but due to their large size, these particles would not be suitable for *in vivo* use.^44^ We chose to use a biocompatible liposomal formulation^45^, which enabled CD33L density to be readily and systematically varied as well as be used *in vivo*. Indeed, CD33L liposomes co-displaying an antigen were successfully used to dampen allergic response in mice through exploiting the inhibitory function of CD33 on mast cells, however, CD33L liposomes in the absence of the allergen had no effect on the allergic response, consistent with the ability of CD33 to antogonize FcεRI signaling, but only when the two receptors are co-ligated.^32^ Modulating the function of microglia in the brain is the next frontier in the treatment of neurodegenerative disease. Clinically used antibodies directly targeting Aβ plaque accumulation have shown some promise but have ultimately not been successful.^46–49^ Thus, augmenting the phagocytic ability of immune cells in the brain may be a better approach. Antibody-based clinical trials are currently underway and while antibodies have long circulatory half-lives, they don’t efficiently cross the blood-brain barrier.^50,51^ Multivalent engagement of Siglecs with glycan ligands offers several advantages over antibodies, including differences in cellular trafficking upon engagement and lower immunogenicity.^24^ CD33 has long been used as a target of leukemia for antibody-drug conjugates, due to the expression of CD33 in AML.^16^ The long-term safety associated with CD33-directed antibody therapies is well established and results demonstrating that mice reconstituted with CD33^-/-^ human immune cells showed normal immune cell function provide further evidence that CD33-engaging therapies will be tolerated.^52^

## Methods

### Reagents and instrumentation

Reagents were purchased from commercial sources as noted and used without additional purification. NHS-pHrodo **4**, anhydrous DMF, and Et_3_N were purchased from Sigma-Aldrich, Canada. NHS-PEG-DSPE **2** and DSPE-PEG-NH_2_ (2000) **5** were purchased from NOF America and Avanti, respectively. CD33L **1** was synthesized through the GlycoNet synthetic core. CDCl_3_ and MeOD_4_ were purchased from Deutero GmbH. Other solvents (analytical and HPLC grade) and reagents were purchased from Aldrich and were used as received. Reactions were monitored by analytical TLC on silica gel 60-F254 (0.25 nm, Silicycle, QC, Canada). Developed TLC plates were visualized under UV lamp (λ max = 254 nm) and charred by heating plates that were dipped in ninhydrin solution in ethanol, and acidified anisaldehyde solution in ethanol. The reaction mixture was purified by silica gel column chromatography (230-400 mesh, Silicyle, Qc, Canada), Sephadex G-100 gel filtration chromatography using CH_2_Cl_2_/MeOH or H_2_O as the elution solvent. NMR experiments were conducted on a Varian 600, or 700 MHz instruments in the Chemistry NMR Facility, University of Alberta. Chemical shifts are reported relative to the deuterated solvent peak or 3-(trimethylsilyl)-propionic-2,2,3,3-d 4 acid sodium salt as an internal standard and are in parts per million (ppm). Coupling constants (*J*) are reported in Hz and apparent multiplicities were described in standard abbreviations as singlet (s), doublet (d), doublet of doublets (dd), triplet (t), broad singlet (bs), or multiplet (m).

### Synthesis of CD33L-PEG-DSPE 3

Chemical reaction scheme is presented in **Supplementary Figure S3**. To a solution of CD33L-NH_2_ **1** (1.86 mg, 1.9 μmol, 1.0 equiv) and NHS-activated DSPE-PEG **2** (5 mg, 1.66 *μ*mol, 1 equiv) in anhydrous DMSO (150 μL, ~10 mM) was added diluted solution of Et_3_N (1.5 equiv.) in anhydrous DMSO and pH of the solution was adjusted between 7.5-8 and the reaction mixture was incubated at room temperature for 6h. An aliquot of the reaction mixture was taken for TLC (CHCl_3_:MeOH:H_2_O = ~75:23:2) analysis. Consumption of the NHS-activated PEG-DSPE and progress of conjugation was monitored by TLC in iodine chamber (to stain CD33L-PEG-DSPE **3** conjugate and NHS-PEG-DSPE **2**), ninhydrin (to stain CD33L amine), and *p*-anisaldehyde (CD33L-PEG-DSPE **3**) staining solution. The solvent was removed and the crude product was loaded to Sephadex G-100 gel filtration column using H_2_O and the crude product was purified using H_2_O as an eluent to afford CD33L-PEG-DSPE **3** as a white powder after lyophilization of fractions having the desired product. Yield: (5.81 mg, 90%), coupling efficiency 70%. ^1^H NMR (700 MHz, MeOD_4_): δ 7.69 (s, 1H), 7.84 (s, 2H), 5.23-5.20 (m, 1H), 5.09 (dd, *J* = 16.1, 48.6 Hz, 2H), 4.40 (dd, *J* = 3.6, 12.6 Hz, 1H), 4.31 (d, *J* = 7.7 HZ, 1H), 4.29 (d, *J* = 7.7 HZ, 1H), 4.19 (d, *J* = 6.3 HZ, 1H), 4.17-4.15 (m, 2H), 4.06-4.02 (m, 1H), 3.97 (t, *J* = 5.6 Hz, 2H), 3.92-3.86 (m, 5H), 3.82-3.80 (m, 3H), 3.73 (t, *J* = 7.7 HZ, 1H), 3.70-3.62 (m, 5H), 3.64 (broad s, 161H), 3.59-3.51 (m, 12H), 3.48-3.46 (m, 1H), 3.42-3.40 (m, 4H), 3.34-3.32 (m, 3H), 3.31 (broad s, 44H), 3.24 (t, *J* = 6.3 Hz, 2H), 3.04 (t, *J* = 6.3 Hz, 1H), 2.34-2.29 (m, 5H), 2.23 (s, 6H), 2.22-2.18 (m, 4H), 2.42-2.22 (m, 2H), 1.89 (t, *J* = 6.8 Hz, 1H), 1.84-1.72 (m, 2H), 1.76-1.73 (m, 2H), 1.60-1.58 (m, 4H), 1.44-1.42 (m, 4H), 1.30 (broad s, 64H), 0.89 (s, 6H); (**Supplementary Figure S4).**The MALDI-TOF-MS spectrum showed the average mass centered at 3.8 kDa and expected average mass was 3.8 kDa (**Supplementary Figure S5)**. The coupling efficiency was determined through assigning aromatic protons signals at 7.69 ppm (s), 7.84 ppm (s) of the phenyl moiety at C9 position of bifunctionally substituted Neu5Ac with terminal methyl groups at 0.89 ppm (s) of the DSPE lipid.

### Synthesis of pHrodo-PEG-DSPE 6

Chemical reaction scheme is presented in **Supplementary Figure S6**.To a solution of NHS-activated pHrodo **4** (1 mg, 1.52 μmol, 1.0 equiv.) and NH_2_-PEG-DSPE **5** (5.3 mg, 1.90 μmol, 1.25 equiv.) in anhydrous DMF (10 mM) was added diluted solution of base Et_3_N in anhydrous to adjust pH of the reaction mixture between 7.5-8 and the solution was stirred for about 2 h at room temperature under dark. Solvent was removed under reduced pressure and the crude was purified by flash silica gel chromatography using gradient elution (MeOH to CH_2_Cl_2_ 1:9 v/v). Yield: (4.90 mg, 88%). Coupling efficiency of **6** with the DSPE-PEG scaffold was determined by ^1^H NMR integration. ^1^H NMR (600 MHz, CDCl_3_): δ 7.54 (d, *J* = 9.6 Hz, 2H),7.45 (s, 1H), 7.10 (s, 1H), 7.01 (d, *J* = 1.8 Hz, 1H), 6.99 (d, *J* = 1.8 Hz, 1H), 6.80 (d, *J* = 1.8 Hz, 1H), 6.67 (d, *J* = 28.2 Hz, 1H), 5.24-5.22 (m, 1H), 4.42-4.22 (m, 1H), 4.20-4.18 (m, 2H), 3.98-3.99 (m, 2H), 3.76-3.68 (m, 2H), 3.64 (broad s, 182H), 3.54-3.51 (m, 2H), 3.48-3.44 (m, 4H), 3.42-3.40 (m, 2H), 3.34 (s, 6H), 2.70 (s, 3H), 2.50 (t, *J* = 6.0 Hz, 1H), 2.31-2.27 (m, 3H), 2.05-2.03 (m, 2H), 1.93-1.91 (m, 2H), 1.27 (broad s, 64H), 1.15 (t, *J* = 6.6 Hz, 3H), 0.89 (s, 6H), 0.85 (s, 6H); (**Supplementary Figure S7**). The MALDI-TOF-MS spectrum showed the average mass centered at 3680 Da and expected average mass was 3680 Da (**Supplementary Figure S8**).

### Synthesis of AF647-PEG-DSPE 7

Compound **7** was prepared by following a procedure described previously.^32^

### Liposome Preparation

Commercially available lipids such as DSPC, Cholesterol, DSPE-PEG were purchased and 10, 5, and 4 mg/mL stock solutions were made respectively. In parallel stock solutions of pHrodo-PEG-DSPE **6** (1mg/mL), AF647-PEG-DSPE **7** (1mg/mL), CD33L-PEG-DSPE **3** (3.2 mg/mL) were prepared. For naked liposomes 57, 38, 4.8, 0.1, and 0.1 mole % of DSPC, Cholesterol, DSPE-PEG, pHrodo-PEG-DSPE **6**, AF647-PEG-DSPE **7** were used respectively. Similar mol % of DSPC, Cholesterol, pHrodo-PEG-DSPE **6**, AF647-PEG-DSPE **7** were used for CD33L liposomes with variable concentration (0.1, 0.33,1, and 3.33 mol %) of CD33L-PEG-DSPE **3** keeping total PEG-DSPE concentration at 5%. Briefly, DSPC, Cholesterol, DSPE-PEG were suspended in chloroform and an appropriate volume of each lipid solution in chloroform was transferred into a glass test tube to reach the desired mol % of each lipid. The solvent was removed under nitrogen gas to form the lipid mixtures. Once all visible chloroform was removed and approximately 100 μL of dimethyl sulfoxide (DMSO) was added to the test tube. CD33L-PEG-DSPE **3,** pHrodo-PEG-DSPE **6**, and 0.1% of DSPE-PEG-A647 **7** in DMSO were then added to the lipid mixture in appropriate proportions to reach the desired mol % of each CD33L. The samples were placed at ≥80°C until completely frozen and excess DMSO was removed via lyophilization overnight and then the dried liposomes were stored at ≥80°C until they were extruded.

Dried lipids were allowed to warm to room temperature and were then hydrated with 1.0 mL of phosphate-buffered saline pH 7.4 (Gibco). The hydrated lipids were then sonicated in a cycle of 1 minute on, 4-5 minutes off until all lipids were uniformly suspended. The lipids were extruded with an 800 nm filter followed by 100 nm filters. The size of the liposomes was then verified by dynamic light scattering (Malvern PanalyticalZetasizer Nano S) to be approximately 120 +-/-20 nm. Liposomes were stored in at 4°C.

### Animal Studies

All mice were on a C57BL/6J genetic background. Transgenic mice expressing hCD33M in the Rosa26 locus were prepared according to previously published methodology.^32^ These mice are widely characterized and tested in our recently published studies.^7,8^ All animals used were maintained in an access-controlled barrier facility under specific-pathogen-free conditions. Studies were approved by the Health Sciences Animal Care and Use Committee of the University of Alberta (AUP00002885).

### Cell lines

WT and CD33^-/-^ U937 used for these studies were generated as a part of another published study.^8^

### Liposome binding studies

U937 cells were grown to a density of ~1 × 10^6^ cells/mL in a T175 flask before the assay, harvested, centrifuged, resuspended in media, and 100,000 cells were added to a 96-well U-bottom plate in 200 μL of media. Cells were centrifuged at 300 rcf for 5 min and the supernatant was discarded. The cell pellet was re-suspended in 50 μL of fresh media and 50 μL of media containing liposomes was added to it. The final concentration of liposomes was 100 μM in each well. The CD33L concentration on the liposomes was 3.33 mol % and the liposome size was 100 nm. These suspensions were incubated for 60 minutes at 37 °C and 4 °C (as a negative control). Following this incubation, 100 μL of media was added to each sample and they were centrifuged at 300 rcf for 5 min. After centrifugation, the supernatant was discarded, and the cell pellet was suspended in flow buffer and further analyzed by flow cytometry. All synthesized liposomes were fluorescently labeled with AF647, which allows us to determine the target specificity of synthesized liposomes to engage the CD33 receptor on the cell surface. The extent of this parameter was determined by assessing the median fluorescence intensity (MFI) of the AF647 fluorescent signal. In each case, cells without liposome treatment were kept to subtract non-specific signals from experimental values.

### Optimization of liposome formulations and their effect on CD33 internalization

U937 cells were grown in a T175 flask and 100,000 cells were added to a 96-well U-bottom plate in 200 μL of media. Cells were centrifuged at 300 rcf for 5 min and cell-pellet was re-suspended in 50 μL of fresh media, and 50 μL of media containing liposome was added to it. These suspensions were incubated for 120 minutes at 37 °C and 4 °C. To best optimize the assay we started with liposome containing different concentrations of ligand (0.1, 0.33, 1, and 3.33 mol %). Furthermore, we varied liposome concentration (1, 10, 100, or 200 μM) to understand how this factor can influence internalization. Finally, we performed a time (10, 30, 60, 120 240, and 360 min) depended-internalization assay with synthesized liposomes. Following the incubation period with the desired liposome, 150 μL of flow buffer was added in each well, and samples were centrifuged at 300 rcf for 5 min. After centrifugation supernatant was discarded and the pellet was stained with 50 μL FITC labeled HIM 3-4 antibody (1:100 dilution from a 1mg/mL stock solution) for 30 minutes at 4 °C. Following this incubation 150 μL of flow buffer was added to each sample and they were centrifuged at 300 rcf for 5 min. After centrifugation, the supernatant was discarded, and the cell pellet was suspended in flow buffer for further analysis by flow cytometry. The extent of decrease of CD33 from cell surface was determined by assessing the MFI of the fluorescent signal observed in FITC channel and using the following formula: Decrease of CD33 from the cell surface (%) = 100X (Cells treated with liposome at 37°C - Cell without liposome treatment 37°C) **/** (Cells treated with liposome at 4°C - Cell without liposome treatment 4°C)

The addition of pHrodo and AF647 fluorophores to liposomes allowed us to determine cellular internalization and binding capability of liposomes respectively. The extent of these two parameters was determined by assessing the MFI of the pHrodo (PE channel) and AF647 fluorescent signal respectively. Each case cells without liposome treatment were kept to determine non-specific signals and treatment at 4°C was treated as a negative control.

### Effect of liposome on cellular phagocytosis

U937 cells were grown in a T175 flask and 100,000 cells were added to a 96-well U-bottom plate in 200 μL of media. Cells were centrifuged at 300 rcf for 5 min and the supernatant was discarded. The cell pellet was re-suspended in 50 μL of fresh media and 50 μL of media containing liposomes was added. The final concentration of liposomes was 100 μM in each well. In parallel, cells were treated with naked liposome which does not contain CD33 ligand. These suspensions were incubated for 60 minutes at 37 °C or 4°C (negative control). Following the incubation period, 100 μL of media was added to each well, and samples were centrifuged at 300 rcf for 5 min. After centrifugation, the supernatant was discarded. The cell pellet was re-suspended in 50 μL of fresh media and 50 μL of fluorescent polystyrene beads (Thermo Fisher) for 30 minutes at 37 °C or 4°C. The final concentration of the polystyrene beads in each well was a 1:100 dilution from a commercially purchased stock solution. Following this incubation, 100 μL of media was added to each sample and they were centrifuged at 300 rcf for 5 minutes. After centrifugation, the supernatant was discarded, and the pellet was suspended in flow buffer for further flow cytometric analysis. The extent of phagocytosis was determined by assessing the percentage of cells taking up at least one bead.

### CD33L-PEG-DSPE insertion into cells

CD33L-PEG-DSPE **3** or PEG-DSPE (1 mM) stock solutions were made in DMSO. To initiate the assay, cells were centrifuged at 300 rcf for 5 minutes and the media was discarded. The cells were resuspended in 50 μL of PBS containing 10 μM CD33L-PEG-DSPE **3** or PEG-DSPE (containing 1% DMSO) and incubated for 1 hr at 37 °C. Cells were centrifuged at 300 rcf for 5 min and the supernatant was discarded. Cells were used in either a phagocytosis assay with polystyrene beads for 30 minutes at 37°C or stained with WT or R119A CD33-Fc according to a published procedure^22^.

### Treatment of primary mouse microglia with CD33L liposome

Adult mice (WT and hCD33M) were euthanized under CO_2_ and their brains were collected and kept in ice-cold RPMI media with 10% FBS, 100 U/mL Penicillin, and 100 μg/ml Streptomycin. Isolated brains were homogenized by 5 ml syringe plungers in media through 40 μm corning filter units under sterile conditions. Homogenized samples were centrifuged at 500 g for 5 min and the pellet was treated with 3 ml of red blood cell lysis buffer (150 mM NH_4_Cl, 9 mM NaHCO_3_, and 0.1 mM EDTA). Following centrifugation at 300 rcf for 5 min, the pellet was dissolved in 3 ml of 30% Percoll (Percoll PLUS, GE Healthcare) and carefully layered on top of 70% Percoll and immediately centrifuged (650 rcf for 20 min). Immune cells were isolated from the border between the two layers, washed (300 rcf, 5 min), and resuspended in media. Isolated microglia from WT and hCD33M mice were mixed and treated with naked liposome or CD33L liposome (100 μM) containing 3.33% ligand for 120 min at 37 °C. After the incubation period 100 μL fresh media was used to wash excess liposomes and cells were centrifuged at 300 rcf for 5 min. Pellet was collected and treated with 200 nM aggregated Aβ or polystyrene beads (1:200 dilution from commercial stock) at 37°C for 30 min. Following this incubation, 100 μL of media was added to each sample and they were centrifuged at 300 rcf for 5 minutes. After centrifugation, the supernatant was discarded, and the pellet was treated with an antibody cocktail containing CD11b (APC/Cy7, clone M1/70, BioLegend), Ly-6G (BV605, clone 1A8, BioLegend), Ly-6C (BV711, clone HK 1.4, BioLegend), Cx3cr1 (PerCP/Cy5.5, clone SA011F11, BioLegend), and F4/80 (BUV395 clone T45-2342, BD Horizon), hCD33 (PE clone WM53 or BV421 clone WM53 BioLegend) at 4°C for 30 min. After the antibody staining, 50 μL flow buffer was added to the cell pellet and centrifuged at 300 rcf for 5 minutes. After centrifugation cell pellet was re-suspended in flow buffer and further analyzed by flow cytometry. To inhibit phagocytosis, cells were pre-treated with 10 μM Cytochalasin-D for 30 min.

### Effect of CD33L liposome in CD33 mutant U937 cell lines

WT and mutant U937 cells were grown in a T175 flask and 100,000 cells were added to a 96-well U-bottom plate in 200 μL of media. Phagocytosis assay was performed with polystyrene beads following the method described earlier. The extent of phagocytosis was determined by assessing the percentage of cells taking up at least one bead. In a separate assay, we also tested depletion of cell surface CD33 by FITC-conjugated HIM 3-4 antibody following the method described earlier.

### Cloning of the hCD33M-FLAG tag fusion protein into the RP172 vector

A 3xFLAG-tag (GATTATAAGGACGACGATGATAAGGATTACAAGGATGATGACGATAAGGACTATAAAGATGACGACGACAAG), containing 5’ AgeI and 3’ AvaI restriction sites and a 3’ stop codon, was synthesized as a gBlocks^®^ Gene Fragment (Integrated DNA Technologies). PCR was used to amplify the sequence and it was cloned into the RP172 vector cut with AgeI. The plasmid was transformed into NEB^®^ Stable competent E. coli cells (New England Biolabs), and isolated plasmids were checked for proper orientation and quality of the FLAG-tag sequence through sanger sequencing. The gene encoding hCD33M, but lacking its natrual stop codon, was subsequently cloned into this vector using the 5’ SphI and 3’ AgeI restriction sites to form the hCD33M-FLAG-tag fusion protein. This was transformed into NEB^®^ Stable competent E. coli cells, and isolated plasmids were checked for quality of the hCD33M-FLAG-tag sequence through sanger sequencing. Lentivirus production and transduction in CD33^-/-^ was carried out the same as a previously published protocol.^8^

### Cell staining and confocal microscopy

Approximately 200,000 CD33^-/-^ U937 cells expressing hCD33M-3xFLAG protein were treated with liposomes (3.3 mol % CD33L or naked) at a final concentration of 100 μM for 45 min. The cells were collected and centrifuged at 300 rcf for 5 minutes and the media was discarded. The cells were washed in PBS and centrifuged. The PBS was discarded, and the cells were resuspended in 100μL of ice cold 3% paraformaldehyde (PFA), followed by 15 min incubation at 4 °C. Cells were centrifuged at 1200 rcf for 5 min, PFA was discarded, and the cells were washed with PBS as described above. For surface staining of CD33, cells were blocked with 100 μL of 5% goat serum for 10 min, followed by incubation with AF488 conjugated anti-CD33 (clone HIM3-4; 1:100 dilution) for 1.5 hr.

To detect internalized hCD33M, we used anti-FLAG antibody (1:100, Sigma). Briefly cells were permeabilized after fixation step by incubation with 100 μL of 5% goat serum containing 0.1% Triton X-100 for 5 min and were further blocked in 5% goat serum for an additional 5 min. The cells were then incubated with anti-FLAG primary antibody for 1.5 hr, followed by centrifugation, and were washed twice with PBS. Cells were incubated with 100 μL of AF488- conjugated anti mouse IgG1 (1:200) for 1 hr, followed by centrifugation and 2 more washed in PBS. Lastly, the cells were incubated with 100μL of Hoechst (1:5000 dilution from a 10 mg/mL stock in PBS) for 15 min and transferred to glass slides. The extra liquid was gently removed, and cells were cover-slipped with antifade permanent mounting medium (TrueVIEW).

Confocal microscopy was performed with the LSM 700 laser scanning confocal microscope (ZEISS). The images were captured at 63X magnification and were analyzed with Zen2.6 Black edition software (ZEISS). A total of 12 cells from each condition were imaged for quantification analysis. For quantification, the sum intensity of CD33 signal in each image was normalized to the Hoechst signal.

### *In vivo* administration of CD33L liposomes via intracerebroventricular (ICV) injection

WT and CD33 transgenic animals were anesthetized with isoflurane inhalation, injected with 2 mg/kg Metacam (analgesic) and immobilized on a stereotaxic device. Stereotaxic marking of the lateral ventricles was performed. Briefly, ICV injections of 1 μl CD33L liposome (20 mM) or naked liposome (20 mM) were performed via syringe needle following craniotomy with pre-optimized coordinates (−1.000 medio-lateral; x, ≥0.300 anterior-posterior, y; and ≥2.600 dorso-ventral relative to bregma, z) to ensure successful injection inside the right ventricle. Injection was performed over 10 min. After injection animals were kept in a recovery incubator for 30 min, and then they were euthanized 4 hr post-injection via CO_2_. Furthermore, their brain samples were collected and primary microglia were isolated according to the methodology described above. After isolation of microglia from both WT and CD33 transgenic mice, a competitive phagocytosis assay was performed with 200 nM fluorescently labeled aggregated Aβ for 30 min at 37°C. After phagocytosis cells were stained with an antibody cocktail and the extent of phagocytosis was measured by flow cytometric methods as mentioned above.

## Acknowledgment

M.S.M. thanks funding through GlycoNet, NSERC, CIHR, and a tier II Canada Research Chair in Chemical Glycoimmunology. A.Bh. thanks GlycoNet for a ATOP fellowship. A.Ba. thanks Alberta Innovates for an undergraduate student research fellowship. AESW was supported by endMS MSSoC and WCHRI graduate scholarships and AV by tier II Canada Research Chair in Neural Stem Cell Biology.

