## Supplementary Information for "Increasing phagocytosis of microglia through targeting CD33 with liposomes displaying glycan ligands"

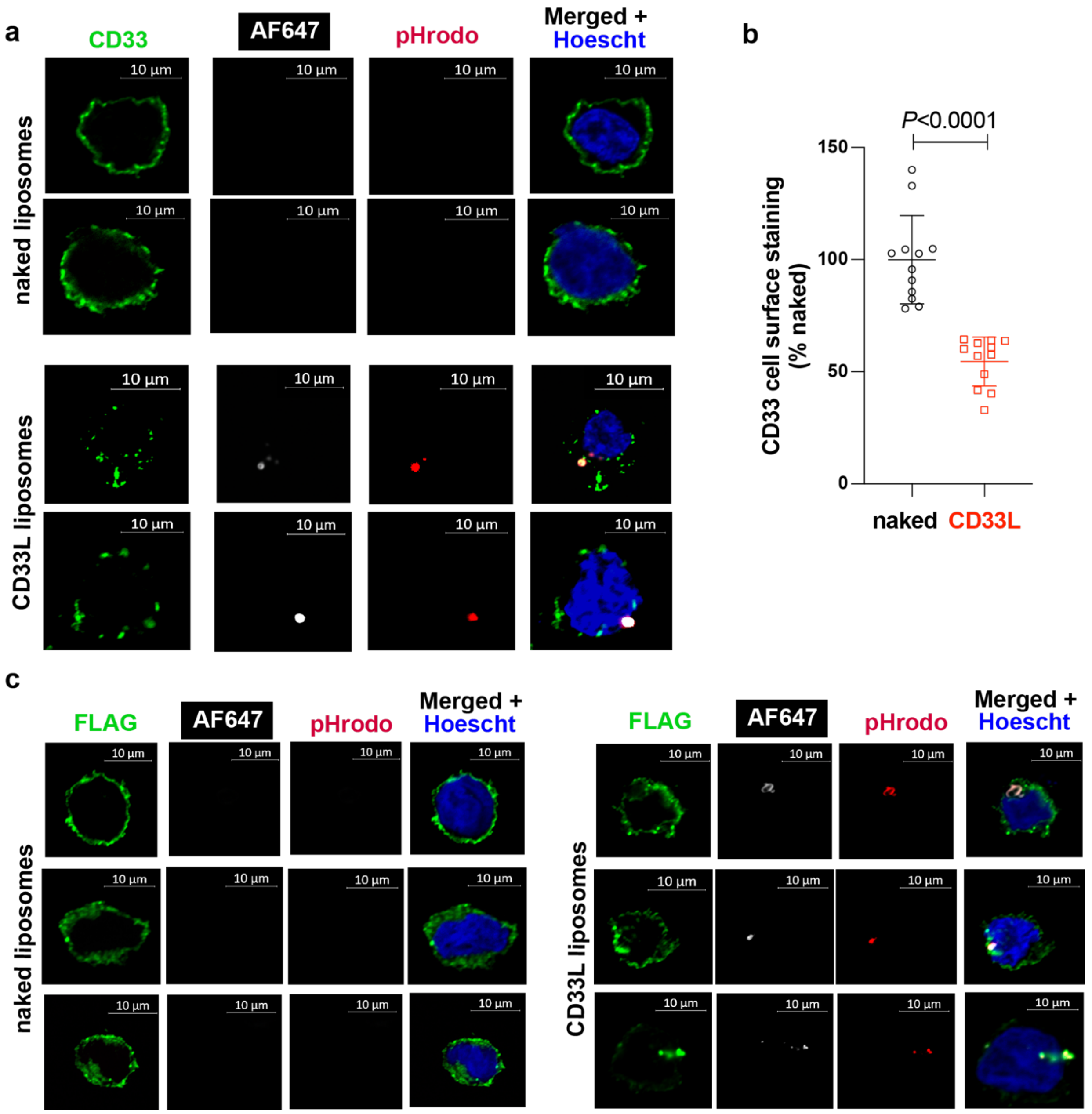


**Supplementary Figure 1: Confocal microscopy demonstrates ligand-induced interanlization of CD33 upon treatment with CD33L liposomes.** (**a**) CD33$^{-/-}$ U937 cells expressing hCD33-3xFlag were stained with anti-CD33 antibody clone HIM3-4 after treatment with naked or CD33L liposomes for 45 minutes. Following incubation, cells were washed and fixed, but not permeabilized. Two representative cells per group are presented. (**b**) Loss of CD33 signal from the cell surface was quantified from 12 cells per each group. Statistical analysis was performed using a two-tailed student's t-test. (**c**) To detect internalization of CD33, another set of cells treated under the same conditions with the liposomes (45 min) were fixed, permeabilized, and stained with anti-FLAG antibody. Three representative images are shown. No fluorescence signal (pHrodo or AF647) coming from internalized liposomes was observed in cells treated with naked liposomes.

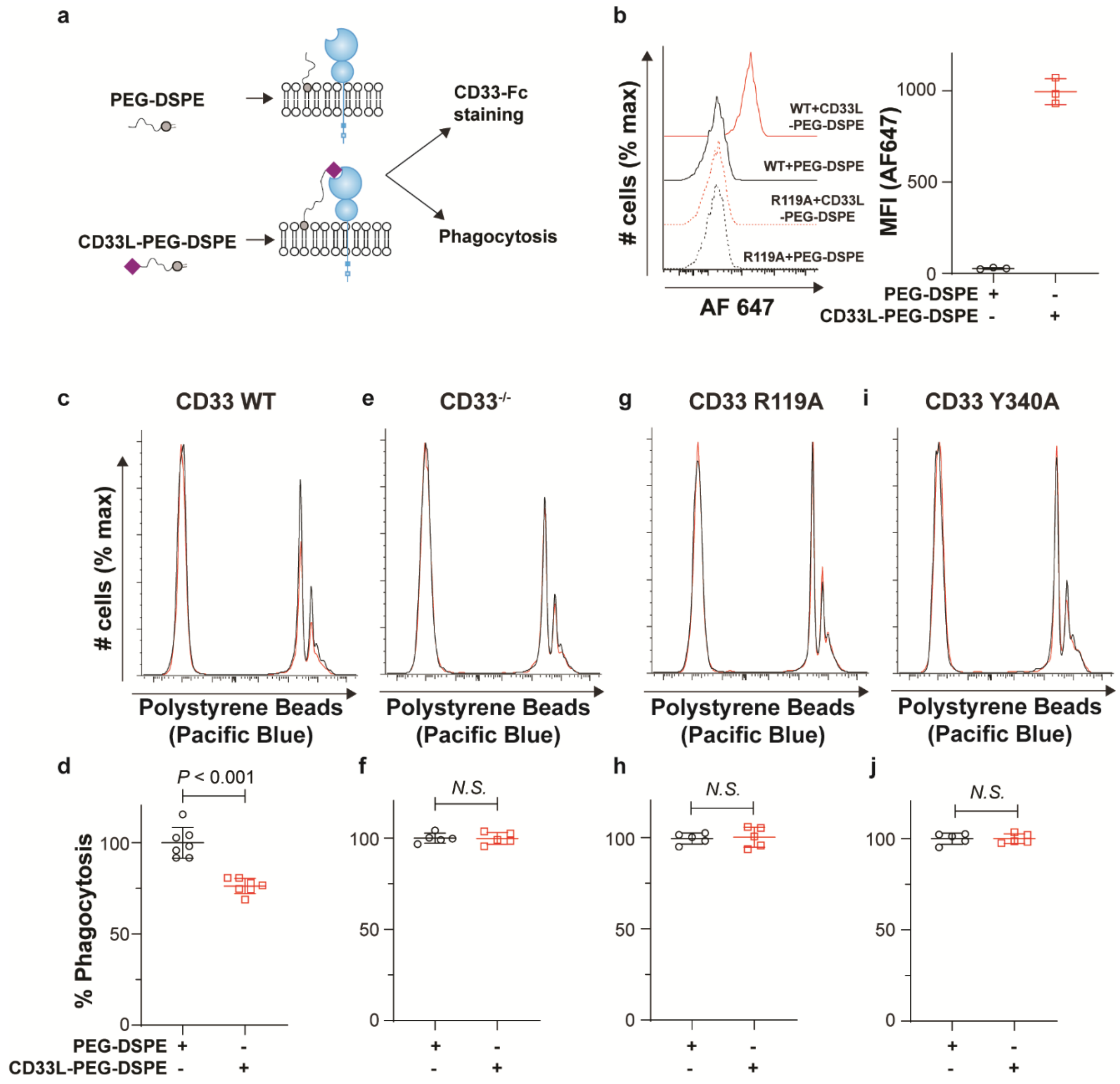


**Supplementary Figure 2. Engaging CD33 with *cis* ligands shows a decrease in phagocytosis.** (**a**) Experimental scheme for insertion of PEG-DSPE or CD33L-PEG-DSPE **3** into WT U937 cells following by probing with CD33-Fc or a phagocytosis assay. (**b**) Staining of U937 cells with WT and R119A CD33-Fc following insertion of PEG-DSPE or CD33L-PEG-DSPE **3**. WT (**c**,**d**), CD33$^{-/-}$ (**e**,**f**), R119A CD33 (**g**,**h**), and Y340A CD33 (**i**,**j**) U937cells were treated with either PEG-DSPE or CD33L-PEG-DSPE **3** prior to performing a phagocytosis assay. Raw flow cytometry plots (**c**,**e**,**g**,**i**) and summary plots of the % phagocytosis (**d**,**f**,**h**,**j**) are shown for each cell type. The assay was replicated five-seven times and Statistical significance was calculated based on an unpaired Student's T-test.

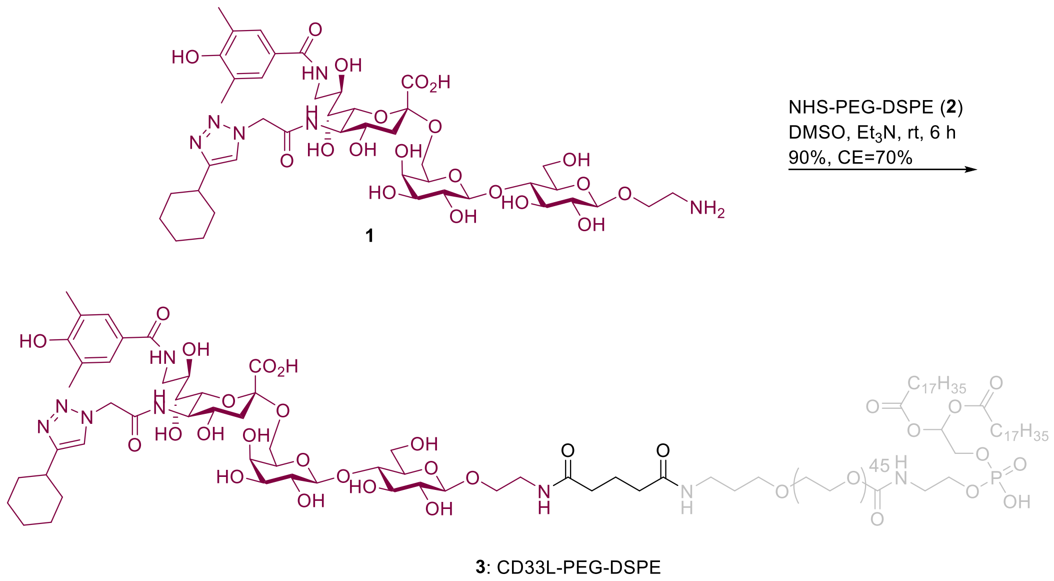


**Supplementary Figure 3.** Reaction synthesis scheme showing experimental conditions required for CD33L-PEG-DSPE **3** production.

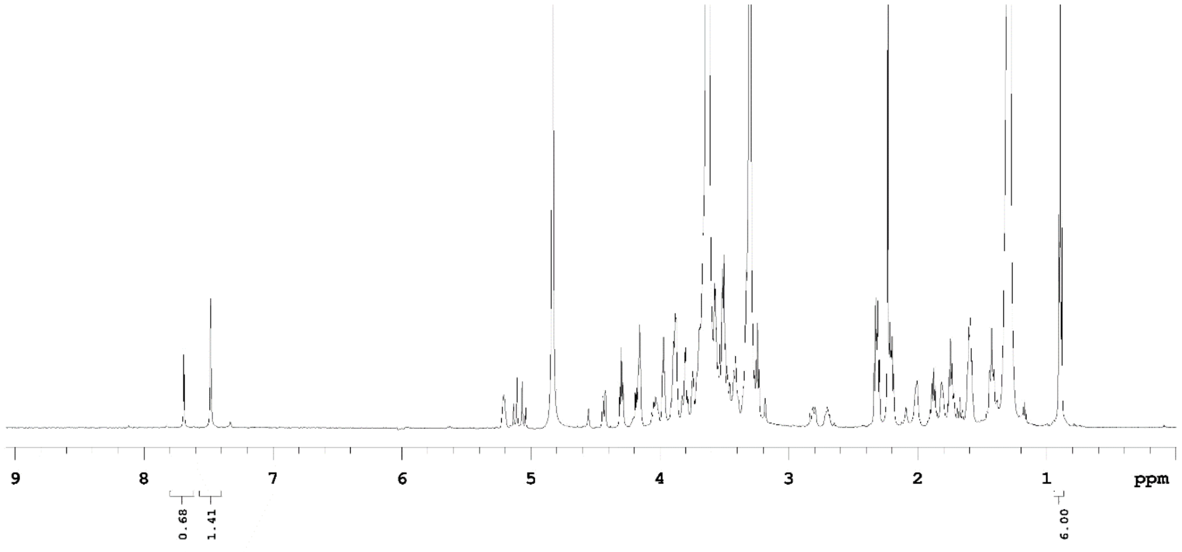


**Supplementary Figure 4.** Purified CD33L-PEG-DSPE **3** is characterized by $^{1}$H NMR (600 MHz, $D_2O$).

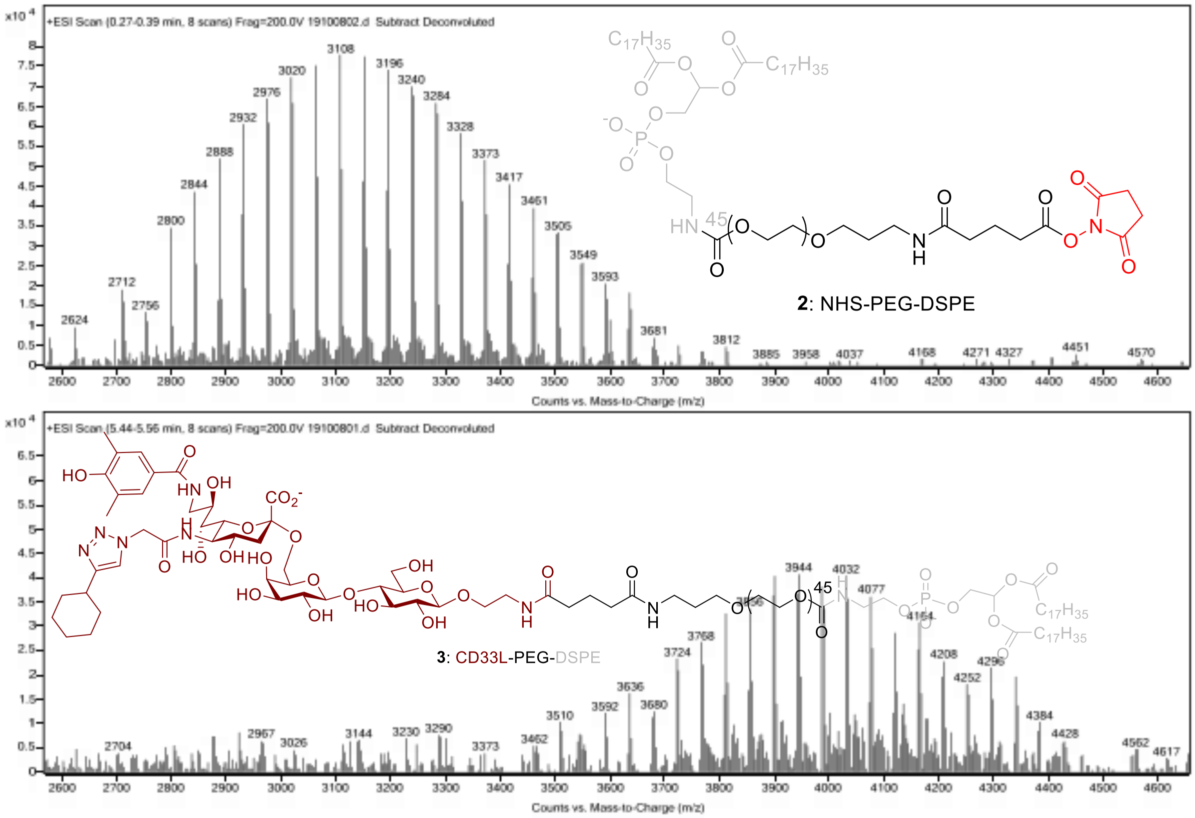


**Supplementary Figure 5.** Molecular weight of CD33L-PEG-DSPE **3** was determined by MALDI-TOF-MS.

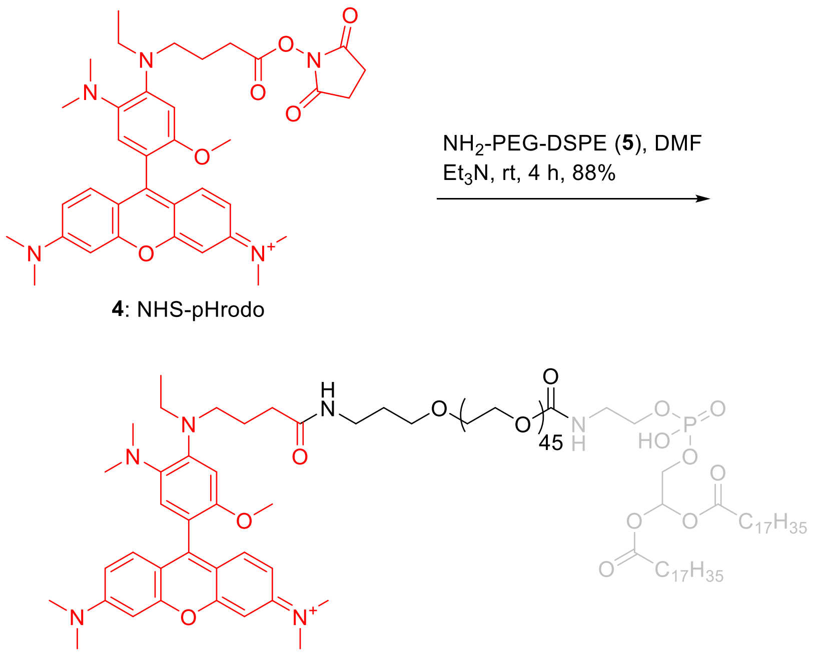


**4**: NHS-pHrodo

**6**: pHrodo-PEG-DSPE

**Supplementary Figure 6**. Reaction synthesis scheme showing experimental conditions required for pHrodo-PEG-DSPE **6** production.

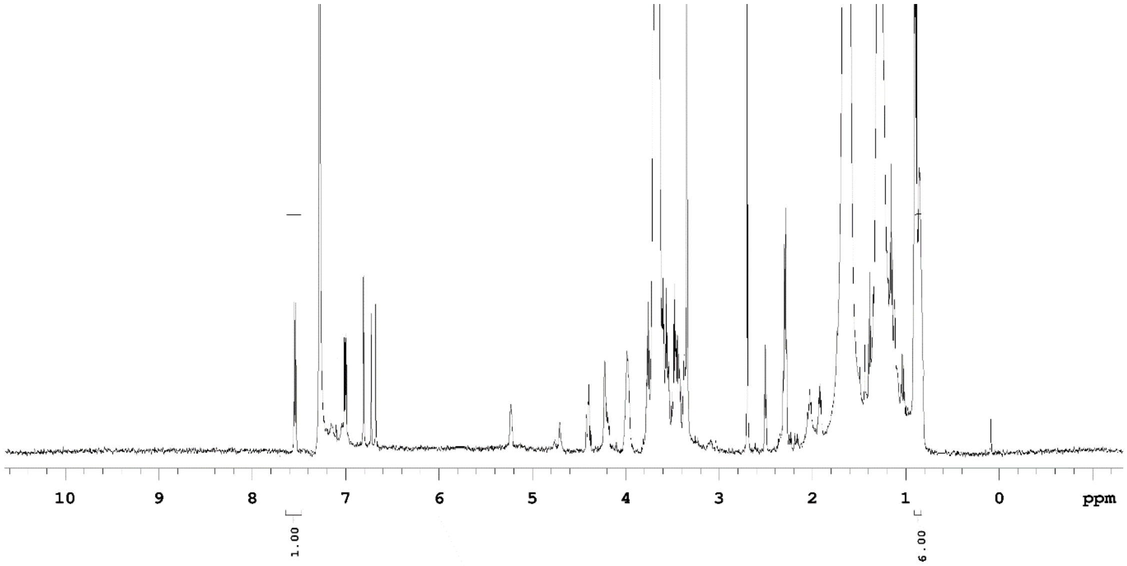


**Supplementary Figure 7.** Purified pHrodo-PEG-DSPE **6** is characterized by $^{1}H$ NMR (600 MHz, $D_2O$).

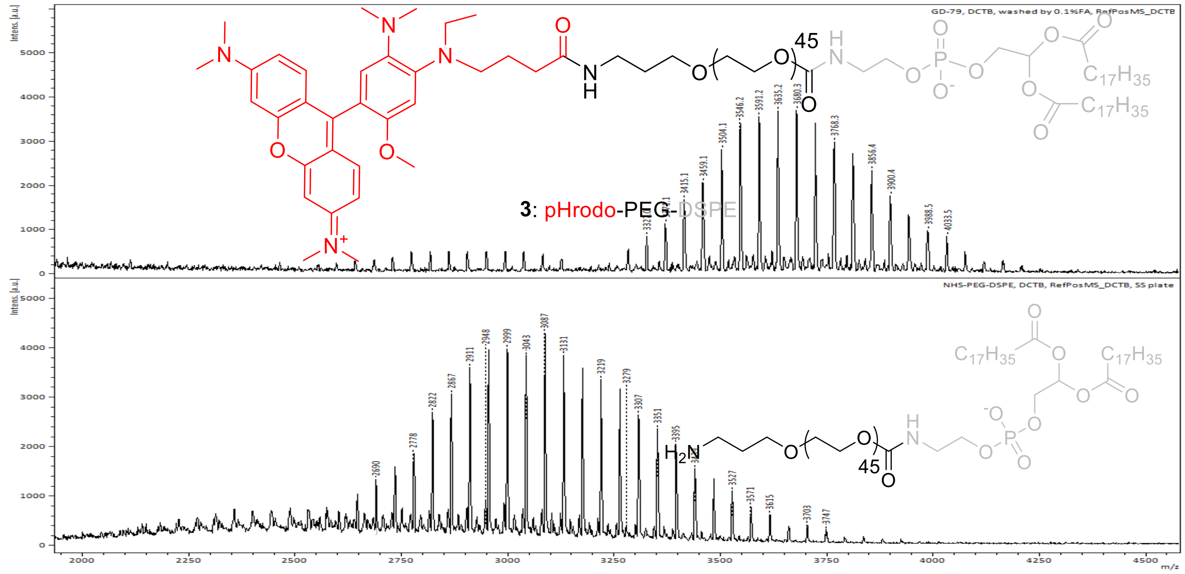


**Supplementary Figure 8.** Molecular weight of pHrodo-PEG-DSPE **6** was determined by MALDI-TOF-MS.
